# Guiding the Refinement of Biochemical Knowledgebases with Ensembles of Metabolic Networks and Machine Learning

**DOI:** 10.1101/460071

**Authors:** Gregory L. Medlock, Jason A. Papin

## Abstract

Mechanistic models are becoming common in biology and medicine. These models are often more generalizable than data-driven models because they explicitly represent biological knowledge, enabling simulation of scenarios that were not used to construct the model. While this generalizability has advantages, it also creates a dilemma: how should model curation efforts be focused to improve model performance? Here, we develop a machine learning-guided solution to this problem for genome-scale metabolic models. We generate an ensemble of candidate models consistent with experimental data, then perform *in silico* ensemble simulations for which improved predictiveness is desired. We apply unsupervised and supervised learning to the simulation output to identify structural variation in ensemble members that maximally influences variance in simulation outcomes across the ensemble. The resulting structural variants are high priority candidates for curation through targeted experimentation. We demonstrate this approach, called **A**utomated **M**etabolic **M**odel **E**nsemble-**D**riven **E**limination of **U**ncertainty with **S**tatistical learning (**AMMEDEUS**), by applying it to 29 bacterial species to identify curation targets that improve gene essentiality predictions. We then compile these curation targets from all 29 species to prioritize refinement of the entire biochemical database used to generate them. AMMEDEUS is a fully automated, scalable, and performance-driven recommendation system that complements human intuition during the curation of hypothesis-driven models and biochemical databases.

**Significance:** Mechanistic computational models, such as metabolic and signaling networks, are becoming common in biology. These models contain a comprehensive representation of components and interactions for a given system, making them generalizable and often more predictive than simpler models. However, their size and connectivity make it difficult to identify which parts of a model need to be changed to improve performance further. Here, we develop a strategy to guide this process and apply it to metabolic models for a set of bacterial species. We use this strategy to identify model components that should be investigated, and demonstrate that it can improve predictive performance. This approach systematically aides the curation of metabolic models, and the databases used to construct them, without relying on the intuition of the curator.

## Introduction

Genome-scale metabolic network reconstructions (GENREs) are knowledgebases describing metabolic capabilities and their biochemical basis for entire organisms. GENREs can be mathematically formalized and combined with numerical representations of biological constraints and objectives to create genome-scale metabolic models (GEMs). These GEMs can be used to predict biological outcomes (e.g., gene essentiality, growth rate) given an environmental context (e.g., metabolite availability) (Oberhardt et al., 2009). GEMs are now used widely for well-studied organisms such as *Escherichia coli* and *Saccharomyces cerevisiae*, but GEMs for most other organisms are much more taxing to create and curate, partially due to the exhaustive and manually-driven steps required (Thiele and Palsson, 2010).

Systems for automatically generating GEMs of sufficient quality for limited purposes have been developed (Henry et al., 2010), but the methods used to further curate GEMs are nearly universally under-reported in the literature. Curation methods for GEMs that take researchers many months to years to develop are often summarized qualitatively with limited description. This is not surprising, given the difficulty in prioritizing areas for curation of network-based, highly connected mechanistic models such as GEMs.

In practice, heuristics are typically used to prioritize curation, such as curating portions of the GEM directly involved in the manipulation of a metabolite, gene, or pathway of known interest. These heuristics, combined with targeted literature searches, allow task-based curation and GEM evaluation, which is increasingly supported in software related to genome-scale metabolic modeling (Lieven et al., 2018; Wang et al., 2018). However, identifying the network components that influence the predictions of interest is not an intuitive process because biological networks are generally highly connected. Gap-filling is an algorithmic approach for identifying reactions to be added to a GEM, or changes to existing reactions, that satisfy imposed constraints on the GEM such as production of a metabolite of interest (Reed et al., 2006). Using gap-filling to guide the curation process is thus limited to helping identify metabolic functions that lead to an experimental phenotype known *a priori*. In other words, gap-filling is a process of fitting a GEM to observed data. This fitting is of tremendous value, but the primary purpose of mechanistic models is to generate *in silico* predictions for behavior in a previously unobserved environment. In order to improve GEM performance for simulation tasks that have no observed experimental equivalent, a curator needs to understand which portions of the GEM affect the output of the simulation.

One way to view this issue is through the lens of a sensitivity analysis, asking how much variation in the parameters of a model will impact a simulation of interest. Such an approach has been developed and applied to dynamic models of biological networks (Babtie et al., 2014), which relies on quantified uncertainty in the structure of a model. Uncertainty quantification has been applied at the level of individual components within a GEM, either by considering the probability of a function being present in a network based on sequence comparisons (Benedict et al., 2014) or by leveraging network structure to more accurately estimate these probabilities (Plata et al., 2012). However, an approach that unifies a probabilistic view of GEM structure with simulations performed with them, which would enable structural sensitivity analysis for GEMs, has not been developed to our knowledge. At a minimum, guiding the curation of a GEM to improve performance on a prospective simulation requires quantifying the uncertainty in the simulation output.

Recently, we developed a framework for the generation of ensembles of GEMs which can be applied to improve predictive performance over that of an individual GEM (Biggs and Papin, 2017). This approach is analogous to the use of ensembles of data-driven models (Dietterich, 2000) or hypothesis-driven models such as signaling networks (Kuepfer et al., 2007), and has been applied to metabolic networks for dynamic modeling as well (Tran et al., 2008). Here, we prioritize curation of GEMs by coupling ensemble modeling with machine learning to take advantage of the uncertainty quantification inherent to ensemble modeling. We call this approach **A**utomated **M**etabolic **M**odel **E**nsemble-**D**riven **E**limination of **U**ncertainty with **S**tatistical learning. (**AMMEDEUS**). One of the central tenets of systems biology is that models represent our hypotheses about how an organism functions. As such, we can use these models to simulate the behavior we expect according to our hypotheses. AMMEDEUS takes advantage of this principle, generating many hypotheses (e.g. an ensemble) and coupling them with machine learning to identify experiments that optimally improve our understanding of a specific behavior for an organism.

## Results

The AMMEDEUS approach is summarized as follows. First, we generate many models that are each consistent with experimental data, forming an ensemble of models (**Figure 1a**). We then perform a set of simulations using the ensemble that are related to a task of interest, such as drug target identification or production of a metabolite of commercial interest (**Figure 1b**). Using the output of these simulations, we perform unsupervised learning to generate phenotypic clusters of models, where clustering is determined by similarity of simulation profiles across the entire set of simulations (**Figure 1c**). We then apply supervised learning to predict simulation cluster membership for each model using the values of variable parameters in that model as input (**Figure 1c**). The relative importance of these model parameters in the supervised learning model indicates the impact that uncertainty in that parameter has on simulation outcomes across the ensemble (**Figure 1d**). In other words, resolving the true state of these parameters will maximally reduce uncertainty in the simulations performed with the ensemble. Here, we apply this approach to the task of reducing uncertainty in predicted gene essentiality for 29 bacterial species (**Figure 1a-d**). We generate an ensemble for each species using previously published growth phenotyping data (Plata et al., 2015), predict the effect of genome-wide single gene knockouts, then apply machine learning as described above. Critically, this process is generalizable to any mechanistic model and simulation task of interest with the correct substitution of machine learning models given changes in the type of simulation output (e.g., continuous vs. discrete, steady-state vs. dynamic).

**Figure 1.**
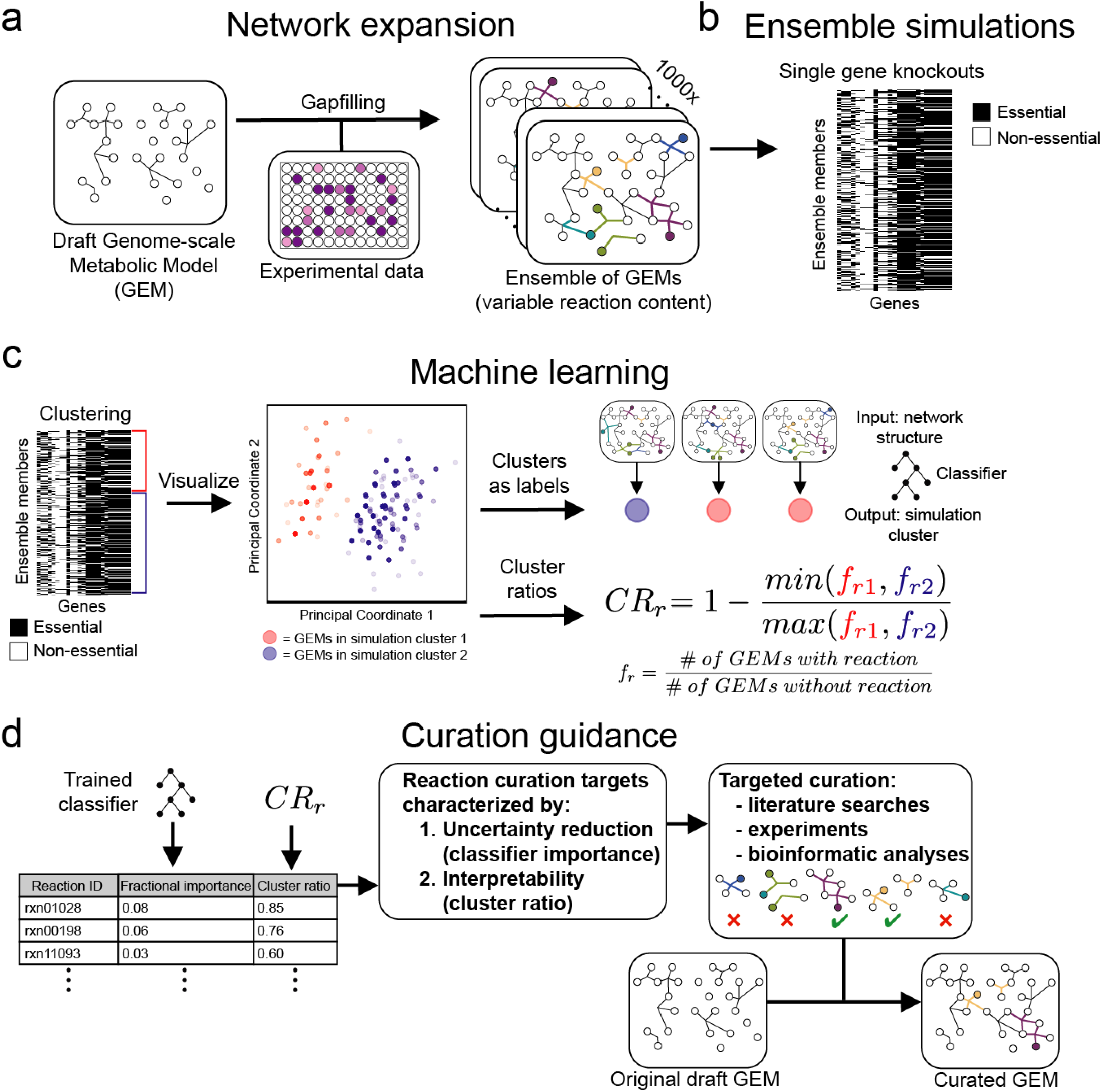
The AMMEDEUS approach guiding curation of genome-scale metabolic models. **a)** A draft GEM is generated using ModelSEED. Algorithmic gapfilling is applied so that the GEM can recapitulate experimental observations, and the process is repeated to identify alternative solutions. These alternative solutions are assembled into an ensemble of GEMs, each of which contains the original content of the draft GEM and a unique set of gapfilled reactions. **b)** Single gene knockouts are performed using the ensemble, in which production of biomass is evaluated when reactions requiring each gene are inactivated. **c-d)** Machine learning approach for identifying curation targets based on ensemble simulations. Unsupervised machine learning is applied to the ensemble simulation results, generating two simulation clusters (cluster 1 [red] and cluster 2 [blue]). Here, principal coordinate analysis is used to visualize the similarity of simulation profiles for all models within an ensemble. Simulation clusters are then used as labels in supervised machine learning, which are predicted using model reaction content as input to a random forest classifier. Curation is prioritized based on the features contributing to classifier performance.

Given our objective of identifying the most impactful experiment or curation effort to improve the quality of a given model, we required ensembles of GEMs that were large enough to saturate the space of unique simulation results (i.e., predicted behavior) and model structures (i.e., hypotheses). We implemented a previously developed iterative gapfilling procedure for generating ensembles of GEMs (Biggs and Papin, 2017). First, each member of an ensemble is generated by iteratively filling gaps in the network to enable *in silico* growth in each of a set of media conditions (**Figure 2a**; see Methods). Alternative solutions are explored by shuffling the order of media conditions used for gapfilling and repeating the process until the ensemble reaches the desired size (**Figure 2b**; see Methods). Using this method, we were able to generate ensembles of around 1000 GEMs for 29 bacterial species (see Methods for descriptions of exceptions).

**Figure 2.**
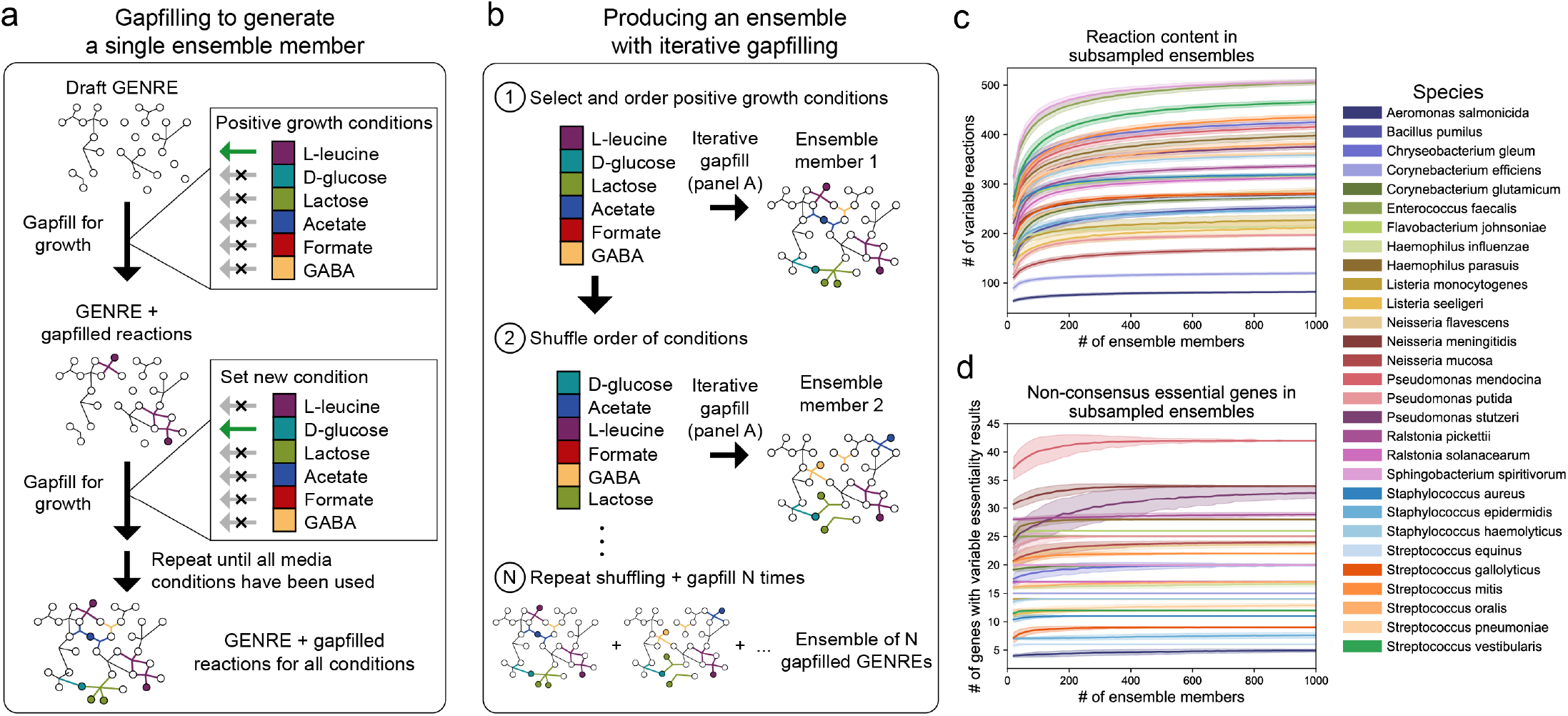
Ensemble generation process and subsampling of ensemble content and simulations to demonstrate adequate sampling of the solution space. **a)** Algorithm for generating a single ensemble member. Given a set of confirmed experimental growth conditions, reactions are added to fill gaps in the GEM required for *in silico* growth in the first condition (See Methods, **Algorithm 1**). The reactions are added, then the process is performed iteratively for the remaining growth conditions until growth is possible in all conditions. **b)** Process for producing an ensemble of GEMs. The algorithm in panel a is performed to generate a single ensemble member, then the order of the media conditions is shuffled and the process is repeated to generate another ensemble member. This process is repeated 1000 times, generating approximately 1000 ensemble members (for some species, duplicate solutions occur and these members are removed. All 29 species had 970-1000 ensemble members) **c)** The variable reaction content in ensemble members as a function of increasing ensemble size. A variable reaction is any reaction that has is variably included across any member of the subsampled group (e.g., it is absent from some members and present in other members, but neither entirely present in nor entirely absent from the ensemble). For each species, the mean number of variable reactions in subsamples of GEMs is shown by the solid line, with the standard deviation shown as light fill of the same color above and below the mean. Subsampling was performed with 1000 draws per subsample size. Ensembles were sampled at intervals of 20 members, e.g., 20, 40, 60… until reaching the size of the entire ensemble. **d)** Variability in gene essentiality simulations within subsamples of ensemble members. Using the same subsampling procedure as in panel c, the number of genes with at least one GEM in the subsample with a simulation outcome different than the rest (e.g., non-consensus) was determined. The mean for each subsample size is shown by the solid line, with the standard deviation shown as light fill of the same color above and below the mean.

To validate that the ensembles we generated represent an adequate sampling of the feasible model space, we first subsampled gap-filled reactions in each ensemble for each species and determined the unique reaction content within each subsample (**Figure 2c**). We found that unique reaction content (e.g., number of unique reactions gap-filled) plateaued or nearly plateaued with ensembles containing as few as 100-200 models, suggesting the ensembles we generated sufficiently saturate the space of unique gap-filled reactions. For gene essentiality simulations, the number of variable predictions (e.g., number of genes for which at least one ensemble member disagrees with another member) plateaued in a similar manner (**Figure 2b**). We also performed subsampling for predictions of growth rate (a common simulation performed with GEMs), which exhibited similar properties of convergence (**Supplemental Figure 1a-b**).

Taken together, these subsampling-based results confirm that ensembles containing 1000 models generated using our reconstruction pipeline sufficiently represent the network structure space (e.g., unique reactions) and prediction space (e.g., essentiality profiles) possible given the input data. This behavior is consistent with our previous work examining the performance of ensembles of GEMs for *Pseudomonas aeruginosa*, in which various aspects of ensemble performance nearly plateaued with only 50 GEMs (Biggs and Papin, 2017). However, in order to ensure that an adequate number of samples are included for downstream machine learning analyses, we maintain the full ensemble of 1000 GEMs for each species in all analyses. In other applications, we suspect that organisms with lower quality GEMs (e.g., more gaps in their metabolic network) or less phenotypic profiling data may require additional sampling to saturate this space. In contrast, species with GEMs containing fewer gaps are likely to require less sampling or an alternative ensemble generation procedure. For example, when attempting to build an ensemble of GEMs for *Bacillus megaterium* using our pipeline, only one unique gapfilling solution could be found. This result is likely due to its large genome size (5.5Mb, 5609 coding sequences) and its extensive genomic and physiological characterization from over 100 years of use in biochemistry research (Eppinger et al., 2011).

Each species’ ensemble contained 19.27 +/− 8.66 genes (mean +/− standard deviation) for which at least one GEM’s prediction of essentiality disagreed with another GEM in the ensemble, representing 3.11 +/− 1.39% of total metabolic gene content. For the unsupervised machine learning portion of AMMEDEUS, we performed *k*-means clustering on the gene essentiality simulations from each species’ ensemble separately. We chose *k* = 2 to generate two clusters for each species, each of which contain GEMs from the ensemble with similar gene essentiality simulation profiles. The results are visualized for all species in this study using principal coordinate analysis (PCoA) in **Figure 3a**. Although we chose *k* = 2 here to illustrate the approach, the separation of models in PCoA space suggests that for many species, determining a larger number of clusters might be advantageous. For example, while *k* = 2 generates two maximally-different simulation clusters, there may be more than two distinct *in silico* phenotypic clusters that represent significant differences in hypothesized model behavior. Accounting for the presence of these smaller clusters may identify important network features that would otherwise only be found through multiple iterations of clustering with *k* = 2 and refinement of the ensemble.

**Figure 3.**
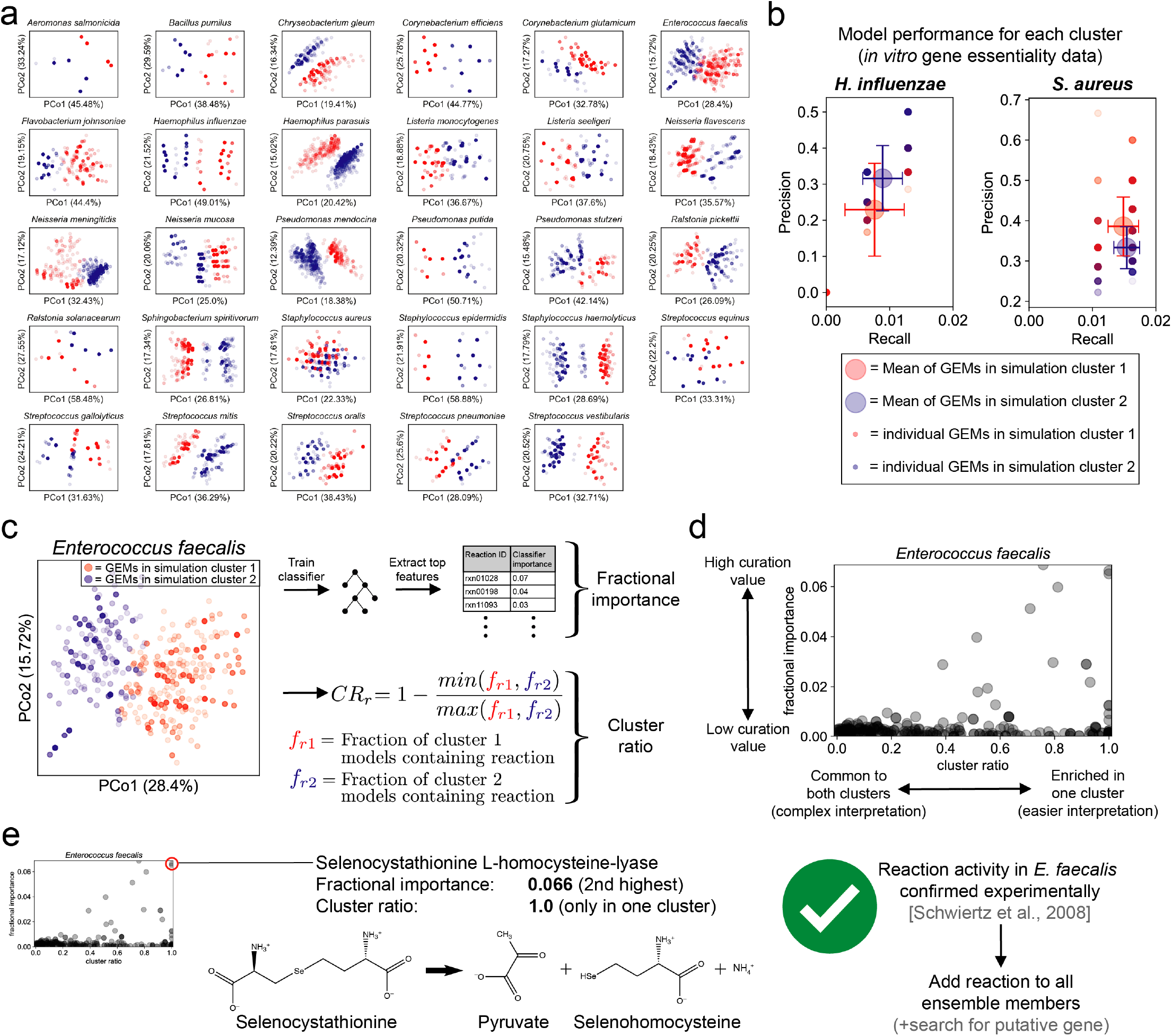
Application of AMMEDEUS to bacterial species. **a)** Ensemble gene essentiality simulations and unsupervised learning for all 29 species. Principal coordinate analysis (PCoA) plots show the similarity between gene essentiality simulation profiles for each ensemble member. Within each PCoA plot, each point represents an ensemble member, colored by cluster membership as determined with *k*-means clustering (*k*=2). PCoA is used solely for visualization; only *k*-means clustering results are used in AMMEDEUS. Percent variance in the pairwise distance matrix explained by each principal coordinate is indicated in parentheses. **b)** Evaluation of performance of GEMs in each simulation cluster compared to genome-wide gene essentiality data. Essentiality datasets from *in vitro* experiments were collected for *Haemophilus influenzae* and *Staphylococcus aureus*. Precision (TP/[TP + FP], TP = true positives, FP = false positives) and recall (TP/[TP+FN], FN = false negatives) were calculated for each ensemble member for each species. Small red and blue circles indicate an individual ensemble member, colored by simulation cluster membership. Large red and blue circles indicate mean behavior for ensemble members from each cluster, and error bars of same color extend above and below the mean by one standard deviation. **c)** Extraction of curation metrics (fractional importance and cluster ratio) for each reaction after the unsupervised learning step. **d)** Example curation guidance plot for *Enterococcus faecalis* **e)** Example of a network feature driving simulation cluster membership. The metabolic activity, selenocystathionine L-homocysteine-lyase, is known to be catalyzed promiscuously by the enzyme cysteine-S-conjugate beta-lyase, which acts on a variety of S- and Se-conjugates. We discovered that this activity has been experimentally verified *in vitro* for *E. faecalis*, but is not incorporated in biochemical databases. Water is excluded from reactants for visualization.

Our approach is focused on prioritizing curation efforts to reduce uncertainty in model simulations. However, whether the parameters we have used result in clusters with differences in predictive performance is unclear. To investigate this question, we evaluated the performance of a subset of ensembles for which experimental genome-wide gene essentiality datasets derived from *in vitro* growth on a rich medium were available. Suitable datasets were identified for *Staphylococcus aureus* (*Chaudhuri et al., 2009*) and *Haemophilus influenzae* (Akerley et al., 2002). Each GEM in the ensemble for each species was evaluated using precision (the ratio of true positives to the sum of true and false positives) and recall (the ratio of true positives to the sum of true positives and false negatives; **Figure 3b**). For both species, ensemble members have variable precision and recall, and simulation cluster membership is associated with a difference in both precision and recall (*p* < 0.0001, Mann-Whitney *U*-test with false discovery rate control via Benjamini Hochberg procedure). We note that the poor precision and recall for all ensemble members is consistent with the performance of other GEMs in predicting gene essentiality, especially when comparing to *in vitro* essentiality datasets that suffer from technical noise and variability (Blazier and Papin, 2019). The critical aspect of these results is that there are biologically meaningful differences in the predictions generated by each cluster. The difference in performance between two clusters suggests that there are meaningful differences in network structure and that assigning two clusters (*k* = 2) is sufficient to capture these differences across an ensemble. Having a meaningful degree of variation between simulation clusters is essential moving forward in AMMEDEUS, as we aim to predict the simulation cluster from the network structure of each ensemble member.

We next sought to identify the reactions that vary across an ensemble that are associated with membership in each cluster. For this objective, we calculated two metrics for each gapfilled reaction in each ensemble. This process is demonstrated for *Enterococcus faecalis* in **Figure 3c-d**. First, we trained a random forest classifier (Breiman, 2001) to predict cluster membership for each GEM from its reaction content. The classifier for every species had an out-of-bag accuracy above 97%, indicating that gene essentiality cluster membership can robustly be predicted from reaction content within the ensembles. To prioritize candidate reactions for curation of each species’ ensemble, we examined the features that contributed the most to classifier performance. We call this first metric the fractional importance of each reaction (called “fractional” because all importances sum to 1 for each species). Second, we developed a metric to represent the enrichment of gapfilled reactions in a single cluster without consideration of classifier performance, which we call the cluster ratio. The cluster ratio (**Figure 3c**) is 1 when a reaction is present in one cluster and not present in any member of the other cluster, and 0 when the reaction is present in an equal number of members in each cluster.

The intent of the cluster ratio is to capture the value of curating reactions that may be lowly abundant throughout an ensemble yet highly enriched in one of the two clusters (e.g., present in 0% of members of one cluster but 20% of members in the second cluster). These reactions may make small contributions to classifier performance due to their low abundance, but curating their presence or absence will reduce the uncertainty in the ensemble of GEMs in a straightforward and interpretable way. This strategy contrasts with reactions with high fractional ratios (i.e., important in the random forest) because the random forest allows for interactions between input variables. As such, curation of individual reactions with high fractional importance may not result in a substantial change in GEM performance; improvements from curating these reactions may be dependent on also curating other reactions that interact with the curated reaction in the trained random forest. Together, the cluster ratio and fractional importance can guide manual curation of a GEM. High cluster ratio reactions represent interpretable “low-hanging fruit” with modest overall value for curation, while high fractional importance reactions represent the highest value curation effort that could be pursued. Reactions with high values for both metrics should be prioritized above all else (**Figure 3d**).

With a list of prioritized reactions for curation in hand, there are multiple approaches that can be taken to curate their presence or absence, including wet lab experiments, targeted bioinformatic analyses, and literature searches. The optimal approach depends on the scientific history of the organism and data availability. A targeted literature search might reveal information that has not been incorporated into genomic or metabolic databases, especially for well-characterized organisms with a large body of literature. For example, in the ensemble for *E. faecalis*, the reaction with the second highest fractional importance and a cluster ratio of 1 (perfectly enriched in one cluster) was selenocystathionine L-homocysteine-lyase, which generates selenohomocysteine, pyruvate, and ammonia from selenocystathionine (**Figure 3e**; reaction IDs: SEED rxn03379 and KEGG R04941). This reaction is catalyzed by cysteine-S-conjugate beta-lyase (Enzyme Commision number 4.4.1.13), which normally catalyzes beta elimination reactions with cysteine sulfur conjugates but is known to act promiscuously on cysteine Se-conjugates (Cooper and Pinto, 2006; Cooper et al., 2011). Cysteine-S-conjugate beta-lyase activity is prevalent in the human gut microbiota, and a study that screened 29 isolates from the gut for Cysteine-S-conjugate beta-lyase and Cysteine-Se-conjugate beta-lyase activity on a variety of conjugates found that *E. faecalis* consistently demonstrated both activities (Schwiertz et al., 2008). Thus, the reaction identified by AMMEDEUS is highly likely to occur in *E. faecalis*, and it should be added to all members of the ensemble to improve the representation of biochemical knowledge as well as reduce uncertainty in the predictions generated by the ensemble. This reaction may be missing appropriate links between genomic and biochemical annotation for a number of reasons: the primary activity is promiscuous (S-conjugates can be one of many compounds), the secondary activity is a less appreciated promiscuous activity (Se-conjugate metabolism by an enzyme primarily known for S-conjugate metabolism), and the primary activity is sparsely annotated in the database used to construct GEMs in this study (PATRIC; contains 45 CDS annotated as “putative cysteine-S-conjugate beta-lyase”, only 7 of which occur outside the *Mycobacteroides* genus). As such, this curation vignette also presents an opportunity for targeted curation of *E. faecalis* annotation in genomic and biochemical databases.

In addition to the single-species curation guidance enabled by AMMEDEUS, the automated nature of the approach allows meta-analyses that span metabolic models for multiple organisms or entire databases. We performed the AMMEDEUS approach for all 29 species in our study. **Figure 4a** shows curation guidance plots for all species, which demonstrate the variability in the distribution of curation target metrics across species. Some species display behavior similar to *E. faecalis*, with many reactions with high fractional importance at intermediate cluster ratio values, indicating complex interactions between reactions of interest (e.g., *Listeria monocytogenes, Listeria seeligeri, Neisseria mucosa*). For these species, reduction of uncertainty in gene essentiality predictions will likely require curation of multiple reactions. Other species have simpler behavior, with a high degree of concordance between cluster ratio and fractional importance for the most important reactions (e.g. *Bacillus pumilis, Haemophilus influenzae, Pseudomonas putida*). For these species, each individual reaction of high importance that is curated will result in a substantial and easily predictable decrease in uncertainty for gene essentiality predictions.

**Figure 4.**
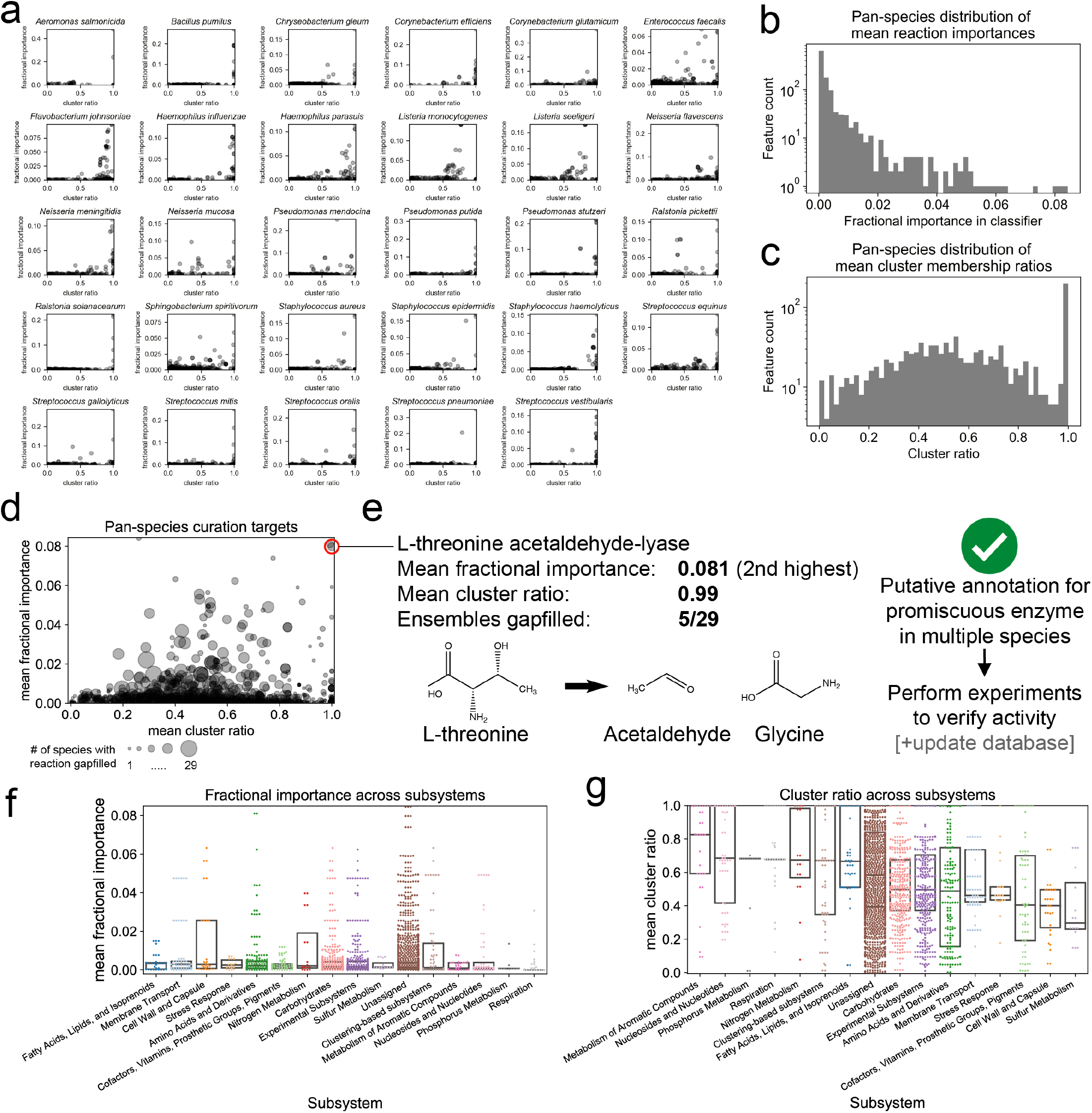
Analysis of AMMEDEUS curation targets for 29 bacterial species. **a)** Curation target plots for all 29 species generated with AMMEDEUS. **b)** Distribution of mean fractional importances across all species. For each gapfilled reaction, fractional importances were averaged across all ensembles in which the reaction was gapfilled; the histogram of these means is shown. **c)** Distribution of mean cluster ratios across all species. Data handled the same as in panel **b. d)** Compiled curation target plot for all species. Size of each point represents the number of species for which the reaction was gapfilled. **e)** L-threonine acetaldehyde-lyase, which had the second highest pan-species mean fractional importance. Species for which the reaction was gapfilled lack an annotated L-threonine aldolase, the primary enzyme known to catalyze this reaction. However, the two species for which this reaction was in the top 10 most important reactions have an annotation for Serine hydroxymethyltransferase, which promiscuously catalyzes this reaction. **f&g)** Mean fractional importance (**f**) and cluster ratio (**g**) across all species by subsystem. Subsystem colors are the same in panels **f** and **g**. Subsystems are ordered by decreasing median from left to right in both panels. Only subsystems with at least 10 gapfilled reactions are shown. Boxplots show median (center line) and extend to the 25th and 75th percentile.

By compiling these curation target metrics across all 29 species, we are able to identify pan-species or database-wide curation targets. For these reactions, improving the accuracy or coverage of gene-protein-reaction associations could greatly improve the performance of GEMs generated with this database for any species. In **Figure 4b**, we show the distribution of mean fractional importances for each reaction used to fill a gap in any ensemble (calculated using the fractional importances only for species for which the reaction was gapfilled). The high-importance tail of this distribution suggests that a small number of reactions have a substantial impact on gene essentiality prediction uncertainty for many species. The same is true of the cluster ratio, for which a large set of reactions have a mean cluster ratio of 1 (e.g., only present in one cluster) across species (**Figure 4c**). The cluster ratio distribution is approximately normal, centered around 0.5, meaning the average behavior for reactions with a cluster ratio not equal to 1 is to be twice as abundant in one cluster than the other (e.g., 1 − ½ = 0.5). These distributional observations are also true when reactions occurring in fewer than 5 species are filtered (**Figures S2a-b**) and when considering the distribution of fractional importances without taking the mean across all species (**Figure S2c**). The distribution of raw cluster ratios (i.e. no mean across species, **Figure S2d**) still has a large set of reactions with a cluster ratio of 1, but has a much larger set of reactions with near 0 cluster ratio (e.g. uniformly distributed across two clusters, 1 − 1/1 = 0). This result suggests that many reactions are evenly distributed between the two clusters for some species, but are enriched in one cluster for at least one other species (resulting in the distribution of means shifting away from 0, as in **Figure 4c**). Taken together, these results suggest that some reactions are of high value across many species (reactions with high mean cluster ratio and/or high mean fractional importance), but these reactions may have minimal or no value for a smaller subset of species.

Individual reactions can be prioritized at the pan-species or database level by taking both cluster ratio, fractional importance, and their frequency across species into account. **Figure 4d** shows the value of each metric for each reaction, as well as the number of species that the reaction was gapfilled for in our analysis. Reactions towards the upper right corner that have large points (i.e., gapfilled for many species) are of highest value from a database curation standpoint. To illustrate a specific example, the reaction with the second highest mean fractional importance is L-threonine acetaldehyde-lyase, which converts L-threonine to glycine and acetaldehyde (**Figure 4e**; reaction IDs: SEED rxn00541 and KEGG R00751; EC 4.1.2.5 and 4.1.2.48). It was gapfilled in 5 out of 29 ensembles, and has a mean cluster ratio of 0.99. This reaction is known to be catalyzed by threonine aldolase (TA) as well as promiscuously by serine hydroxymethyltransferase (SHMT; generally encoded by *glyA*) in bacteria (Chaves et al., 2002). For two species in this study, *Corynebacterium efficiens* and *Haemophilus parasuis*, this reaction is amongst the 10 reactions with the highest fractional importance. TA activity is known to occur in *Corynebacterium glutamicum*, a close relative of *C. efficiens*, but it is not known whether the activity is due to TA or SHMT (*Simic et al., 2002*). Interestingly, the genomes for *C. efficiens* YS314 and *H. parasius* SH0165 (both used in this study as representative genomes) contain a putative SHMT encoded by glyA, but no putative TA. For these species, a simple experiment with crude extracts to verify TA activity, as performed previously for *C. glutamicum*, could verify that the metabolic activity occurs either through promiscuous SHMT activity or an orphan enzyme (Simic et al., 2002). A more systematic set of experiments utilizing glyA mutants for these species and a handful of others could identify the degree of promiscuous TA activity by SHMT to properly propagate annotations to other species within databases. In addition to the inherent value in improving the quality of biochemical databases through this targeted investigation, AMMADEUS shows that this specific investigation would substantially decrease uncertainty in gene essentiality predictions for a broad selection of bacterial species.

Based on this pan-species analysis, we next asked whether reactions within specific subsystems were contributing more to prediction uncertainty than other pathways. Reactions assigned to “Respiration” had a lower mean fractional importance than reactions assigned to any other pathway except for “Phosphorus metabolism” (**Figure 4f**, Kruskall-Wallis test with posthoc pairwise Dunn’s test and Bonferroni multiple testing correction). Given the key role of respiration in energy generation and it’s well-characterized structure, it is no surprise that reactions directly involved in respiration do not significantly contribute to prediction uncertainty for GEMs. Few other differences in mean fractional importances across pathways exist (see **Supplemental Table S2**). The same analysis, instead performed on the mean cluster ratio of reactions assigned to each subsystem, yielded far more differences between subsystems (**Figure 4g, Supplemental Table S3**). “Metabolism of Aromatic Compounds” and “Nucleosides and Nucleotides” had particularly high mean cluster ratios, suggesting that curating individual reactions within those subsystems should reduce prediction uncertainty without dependence on curating larger pathways. “Respiration” also had a high cluster ratio, in contrast to its low mean fractional importance. One potential explanation for this is that the small number of reactions involved in respiration tend to be essential, but they also have overlapping roles for generating key metabolites such as pyruvate, L-lactate, and acetyl-CoA, and thus may be redundant for some GEMs. Reactions involved in respiration also have redundant electron carriers, such as ubiquinone and menaquinone, so preferential addition of each of the two reactions to different clusters could result in a high cluster ratio for each reaction without any impact on gene essentiality simulations. Other subsystems had mean cluster ratios centered closer to 0.5, similar to the mean value in the normal portion of the distribution in **Figure 4c**. Together, the subsystem-specific fractional importance and cluster ratio behavior suggests that focusing on individual reactions in database-wide curation will have greater value than focusing on subsystems. Thus, in practice, modelers that aim to improve their GEM with respect to a broad set of simulations (e.g. genome-wide gene essentiality) should focus curation on key reactions that are distributed across the network rather than curation of specific predefined subsystems. AMMEDEUS provides a systematic way to identify these reactions for individual organisms and entire biochemistry databases.

## Discussion

The analysis we performed demonstrates just one possible path towards the goal of reducing uncertainty in our understanding of biochemical networks within the AMMEDEUS framework. Changes to the process can be rationalized for new goals; for example, we previously demonstrated that introducing random weights on inclusion of each reaction during algorithmic gapfilling can generate more diverse ensembles (Biggs and Papin, 2017). If none of the ensemble members generated by our pipeline adequately represented metabolism for an organism (e.g., their gene essentiality simulation results were vastly different than experimental observations), we could introduce such random variance to increase the likelihood of generating some ensemble members that reflect biological reality. Such an approach may be necessary for organisms with metabolic repertoires differing substantially from those represented in popular biochemical databases (e.g., gut microbes, intracellular parasites). Inclusion of methods for proposing novel hypothetical enzymatic function could complement our approach for such organisms (Hatzimanikatis et al., 2005; Jeffryes et al., 2015).

AMMEDEUS can be immediately extended to other simulations performed using GEMs with small adjustments to the machine learning models applied. For example, rather than gene essentiality, we may be interested in improving growth rate predictions across many media conditions. In this case, we would perform ensemble flux balance analysis in each condition to predict growth rates (Biggs and Papin, 2017), then apply an unsupervised machine learning algorithm suited to continuous data, such as principal component analysis (PCA). In this setting, each sample would be a vector of growth rates generated by a single ensemble member, and the loadings in PCA would describe variance in predicted growth rates, and each sample (ensemble member) would have a score for each principal component. In the supervised learning step, we would apply regression to predict the scores (e.g. predict the value of the first principal component [PC1] for each sample) using the presence or absence of gap-filled reactions as the regressor input. The feature importances in this regressor would be equivalent to the fractional importances in the random forest classifier we use in the implementation of AMMEDEUS in this study. To calculate an equivalent to the cluster ratio, the same equation could be used with *f*_1_ and *f*_2_ replaced with the absolute value of the average of PC1 for ensemble members with and without the reaction, respectively. This hypothetical shift in curation goals, and the simple swapping of machine learning models required, demonstrates the modular nature of AMMEDEUS.

Our approach builds on work in other disciplines in which uncertainty quantification and reduction are applied to understand or improve the behavior of domain-specific models. For example, in petroleum engineering, an ensemble-based approach is used to derive value of information (VOI) estimates for resolving parameter values in models of oil reservoir management (He et al., 2018). In this setting, a company may be interested in performing the experiment or analysis needed to improve their certainty in a model of profit gain or risk. With AMMEDEUS, we effectively derive VOI estimates for resolving reaction presence or absence, where value is determined by the degree of uncertainty reduction for predictions of interest. Taking a VOI approach for biological discovery and to improve the models used in various facets of biotechnology could help automate workflows and substantially reduce costs by prioritizing experiments. Machine learning methods have great utility towards this goal, since they can be applied to any variety of mechanistic model structures and simulation outputs, removing the need to derive analytical solutions for VOI estimates for every new scenario. As the diversity and depth of organisms that mechanistic models such as GEMs are being constructed for increases, such approaches will be vital to continue to improve their quality and predictiveness (Magnúsdóttir et al., 2017; Monk et al., 2014).

## Methods

### Organism selection

Organisms with available growth phenotype data were extracted from Plata, *et al*. (Plata et al., 2015). To identify a representative genome for each species, we queried the PATRIC database (Wattam et al., 2017) with the genus and species name for all organisms in the study, then selected a single genome from PATRIC based on decision criteria described as follows. When a reference genome was assigned for the species, the genome identifier for the reference genome was chosen. If no reference genome was available, a genome listed as “representative” was chosen. When multiple genomes with the “representative” status were available, we chose the first genome listed. If a selected representative genome contained more than 10 contigs, a representative genome with fewer contigs was chosen. Strain identifiers were not provided in the study from which data was drawn, so these selection criteria were developed to select the highest-quality genome available for the species in the study. Selected genome identifiers are available in Supplemental Table S1.

Organism selection was further refined by only including those from Plata, *et al*. (Plata et al., 2015) which grew in at least 10 of the single-carbon source Biolog conditions. The experimental growth threshold originally used in the paper from which data were drawn was used (>10 colorimetric units of tetrazolium dye reduction; originally scaled between 0 and 100 based on positive [100 units] and negative [0 units] controls). This choice was made with the recognition that the tetrazolium dye measures redox activity and not actual biomass production; for the purpose of our study, we assume that detectable redox activity above 10 relative units would require biomass production. After this initial selection step, *Brachybacterium faecium* and *Gordonia bronchialis* were also removed from the analysis because no solutions existed to enable biomass production using the universal reaction bag for either species. *Bacillus megaterium* was excluded because only one gapfill solution was found across all gapfilling cycles. Similarly, *Stenotrophomonas maltophilia* was excluded because only two unique gapfill solutions were found. In total, the full analysis pipeline was applied to 29 species.

### Generation of draft genome-scale metabolic models

Draft-quality genome-scale metabolic models (GEMs) were generated using the ModelSEED reconstruction pipeline (Henry et al., 2010) accessed through PATRIC in August 2018 (Wattam et al., 2017). PATRIC servers were queried to generate GEMs formatted for use in cobrapy (Ebrahim et al., 2013) using the Mackinac package (Mundy et al., 2017).

### Representative media

The base medium for biolog conditions was derived from the ModelSEED media compositions for biolog plates. Flux variability analysis was used to identify metabolites which had essential uptake reactions in all complete media-gapfilled reconstructions from PATRIC. Based on this analysis, we added Heme and H2SO3 to the base biolog composition used *in silico* (i.e., uptake of heme and H2SO3 was allowed in all conditions). For each single carbon source, appropriate identifiers were found in the ModelSEED database. For metabolites with ambiguous chemical identities (e.g., metabolites that Biolog does not provide isomer composition for, such as D-galactose), only one isomer was selected from ModelSEED to represent the condition. Carbon sources that are complex mixtures of metabolites (gelatin) or polymers (pectin) were excluded from analyses.

### Algorithmic gapfilling

Each individual gapfilling step, corresponding to enabling biomass production on a single media source, was performed using the following algorithm adapted from our previous work (Biggs and Papin, 2017). This algorithm is in essence the same as parsimonious flux balance analysis (pFBA, (Lewis et al., 2010)), except that the parsimonious assumption of minimization of the sum of all fluxes is only applied to reactions from a universal reaction bag that are activated to allow flux through the network.

Algorithm 1: pFBA-based gapfilling

Min Σ(*abs*(*y_j_*)) *for j* ∈ [0,1, … # *universal reactions*], subject to:

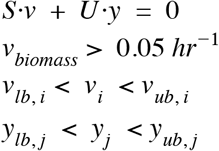

Where *S* is the stoichiometric matrix representing the model to be gapfilled, *v* is the vector of fluxes through reactions in *S, U* is the stoichiometric matrix representing the reaction database from which reactions are activated to fill gaps, *y* is the vector of fluxes through reactions in *U, v_biomass_* is flux through the biomass reaction, *v_lb_* and *v_ub_* are lower and upper bounds of flux through reactions in the original model, respectively, and *y_lb_* and *y_ub_* are lower and upper bounds of flux through reactions in the reaction database *U*.

The formulation is identical to the original formulation of pFBA, except for four key differences. First, we only require an arbitrarily low amount of flux through biomass, rather than the maximum amount of biomass, meant to represent a binary growth condition. Second, we introduce a universal reaction bag (*U*) and associated flux variables for each reaction in U (*y*). Third, only fluxes through reactions in *U* are penalized; fluxes through reactions in the model being gapfilled (*S*) are not penalized. Fourth, rather than explicitly splitting all reactions into irreversible reactions, we take advantage of solver-level interfaces implemented in cobrapy through the optlang package (Jensen and Cardoso, 2016) that allow introduction of absolute values into the objective (this is done out of convenience in our implementation; this aspect of the problem formulation is identical to the same aspect in pFBA at the solver level) (Jensen and Cardoso, 2016). As in Biggs et al. (Biggs and Papin, 2017), the solution to this optimization problem activates reactions in the universal reaction bag with the minimum sum of fluxes necessary to enable flux through the biomass reaction in a given condition.

### Generating ensembles from gapfill solutions

For each organism, the entire algorithm for generating an ensemble is as follows:

~~~
For i in number_of_ensemble_members:
      Randomly order a selection of J media conditions
      For a single condition j in J:
         1. Set model bounds to represent media condition
         2. Optimize using pFBA-based gapfilling
         3. Add activated reactions/remove flux
            minimization penalty for those reactions
      Store solution from this iteration
Create an ensemble where each member contains the set of
      reactions added over an iteration through all media conditions.
~~~

We performed this procedure for 1000 cycles for each species (i.e., `number_of_ensemble_members = 1000`). All species included in the study grew in at least 10 *in vitro* single carbon source media conditions (i.e., j contained at least 10 conditions); for each species, all positive growth conditions were used to gapfill during each cycle. After removing duplicate gapfill solutions, all species included for further analyses had 970-1000 members in their ensemble (species not considered after this point are detailed in *Organism selection*).

### Ensemble Flux Balance Analysis and Ensemble Gene Essentiality

Ensemble flux balance analysis and ensemble gene essentiality screens were performed using Medusa v0.1.2 (Medlock and Papin, 2019) and cobrapy v0.13 (Ebrahim et al., 2013). The GNU linear programming kit (GLPK) was used as the numerical solver in all cases. For all simulations, rich medium was used (1,000 mmol/gram dry weight*hr uptake allowed for all metabolites with a transport reaction; commonly referred to as “complete medium”). An arbitrarily low cutoff for flux through biomass in gene essentiality screens was used (1E-6 units of biomass/hr), but varying this quantity between 1E-10 and 1E-3 did not substantially affect essentiality results.

### Subsampling of ensemble features and predictions

For all subsampling performed, 1,000 random draws were made with replacement at each subsample ensemble size. Ensemble sizes for each subsampled population ranged from 20 to 1,000, with subsampling performed in intervals of 20 members (i.e., 20, 40, 60 … 1,000 members). When the subsample size exceeded the actual ensemble size (e.g., some species had slightly less than 1,000 members), all ensemble members were subsampled.

### Clustering of ensemble gene essentiality predictions and prediction of clusters

Prior to clustering of gene essentiality predictions, genes with perfectly correlated predictions across an ensemble were collapsed to a single variable (i.e., if gene 1 always has the same essential/nonessential prediction as gene 2, they are lumped as a single variable). Without this aggregation, these perfectly correlated features heavily biased *k*-means clustering resulting in unbalanced clusters with ~90% of ensemble members in a single cluster. After aggregation of perfectly correlated genes, ensemble gene essentiality predictions were clustered into two clusters using *k*-means clustering as implemented in the KMeans class of scikit-learn v0.19.2 (Pedregosa et al., 2011) (max iterations=300, convergence tolerance=1E-4, Elkan’s(Elkan, 2003) algorithm). Gene essentiality predictions were converted to binary data (essential or nonessential) using a cutoff of flux through biomass of 1 E-6 mmol/(gDW*hr). Random forest classification was performed to predict cluster membership using active features in each ensemble member (e.g., presence or absence of a reaction was assigned as True or False in the input, respectively) (Breiman, 2001). The RandomForestClassifier class from scikit-learn v0.19.2 was used (500 trees, quality of splits determined with the gini criterion, no max depth, minimum of 2 samples per split, minimum of 1 sample per leaf, sqrt(number of features) searched at each split, training samples determined for each tree via bootstrap selection with replacement). The default metric in scikit-learn’s RandomForestClassifier for determining feature importance, the mean decrease in node purity, was used to calculate feature importance in this study (Gordon, 1984).

### Visualization of gene essentiality clusters

Principal coordinate analysis (PCoA) (Gower, 1966) was used to visualize ensemble gene essentiality results. PCoA as implemented in scikit-bio v0.5.4 (https://github.com/biocore/scikit-bio) was performed using the hamming distance (Hamming, 1950) to compute the pairwise distance matrix.

### Gene essentiality datasets

Gene essentiality datasets were identified for species in this study from the Online Database of Gene Essentiality (OGEE, (Chen et al., 2017)). In cases where multiple datasets were available for a given species, the dataset generated using the same strain of the species selected for GENRE reconstruction was selected. If multiple datasets still existed for a species, a single dataset was chosen based on media richness (e.g., more complex media were selected over simpler media). We excluded the essentiality dataset for *Streptococcus pneumoniae* because the total set of screened genes was not included (Song et al., 2005). In brief, the authors developed a kanamycin insertion cassette targeted for 693 genes that were selected based on having >40% amino acid sequence identity with a set of well-studied organisms. The authors reported the identity of only the essential genes, so non-essential genes that would be in the dataset could not be included in our set of predictions. Based on these selection criteria and limitations, we selected datasets from OGEE for *Staphylococcus aureus* (*Chaudhuri et al., 2009*) and *Haemophilus influenzae* (Akerley et al., 2002).

### Subsystem analysis

Subsystem assignment for reactions in ModelSEED were obtained from the ModelSEED biochemistry repository in April 2019 (https://github.com/ModelSEED/ModelSEEDDatabase/blob/dev/Biochemistry/Pathways/ModelSEED_Subsystems.tsv). The highest level subsystem assignment, “class”, was used. For reactions with multiple subsystem assignments at this level, the reaction was considered as a separate observation belonging to both subsystems with the same mean fractional importance and mean cluster ratio (e.g., a reaction belonging to two subsystems is an independent observation for each subsystem in **Figures 4f&g**). To test for differences amongst subsystems, we performed a Kruskal-Wallis test with post-hoc pairwise Dunn’s tests with Bonferroni multiple testing correction using SciPy version 1.1.0 (Kruskal-Wallis) and scikit-posthocs version 0.6.1 (Dunn’s test with Bonferroni correction) (Jones et al., 2016; Terpilowski, 2019).

### Data and analysis availability

All data, analysis scripts, results, and models generated are available at https://github.com/gregmedlock/ssl_ensembles and will be archived on Zenodo upon acceptance for publication after peer review.

## Acknowledgements

We acknowledge funding from the National Institutes of Health R01GM108501, R01AT010253, T32LM012416, the Thomas F. and Kate Miller Jeffress Memorial Trust, and a Wagner predoctoral fellowship to GLM. We thank Matthew Biggs for thoughtful discussion related to the manuscript and Maureen Carey for helpful comments on drafts.

## Author Contributions

Conceptualization, G.L.M and J.P; Data Curation, G.L.M; Formal Analysis, G.L.M; Investigation, G.L.M; Methodology, G.L.M; Software, G.L.M; Validation, G.L.M; Visualization, G.L.M; Writing - original draft, G.L.M; Writing - Review & Editing, G.L.M and J.P; Funding Acquisition, G.L.M and J.P; Project Administration, J.P; Resources, J.P; Supervision, J.P.

## Supplemental Figures

**Figure S1.**
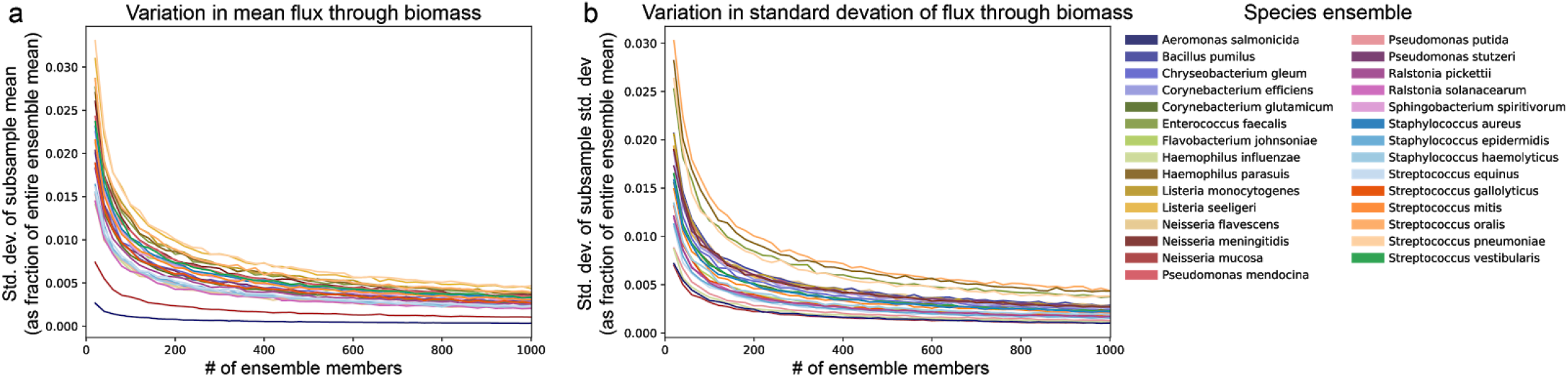
Subsampled ensemble behavior for predictions of biomass production. We simulated biomass production in a rich medium across the entire ensemble and subsampled these results at varying ensemble sizes. **a)** Standard deviation of the mean flux through biomass from each subsample and **b)** standard deviation of the standard deviation of flux through biomass in each subsample. For both quantities (variance of the mean of each subsample and variance of the variance of each subsample), simulations plateau before inclusion of all 1000 ensemble members. Values on the y axis are normalized by dividing by the mean flux through biomass for the entire ensemble.

**Figure S2.**
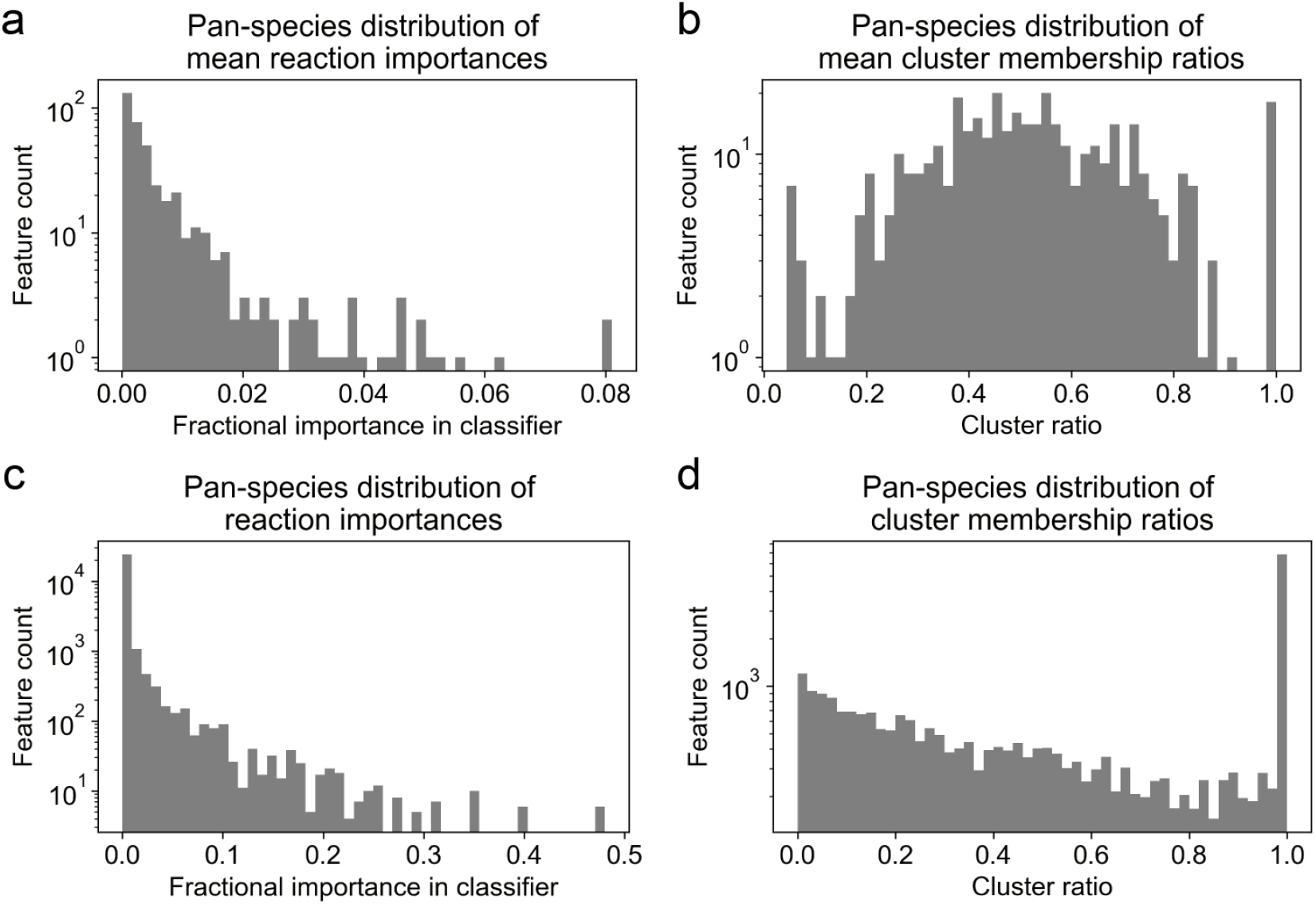
Distribution of fractional importances and cluster ratios. **a)** Distribution of mean fractional importances for reactions gap-filled in at least 5 ensembles. Identical to **Figure 4b** other than filtering step. **b)** Distribution of mean cluster ratios for reactions gap-filled in at least 5 ensembles. Identical to **Figure 4c** other than filtering step. **c)** Distribution of reaction importances across all species. Identical to **Figure 4b** except the mean is not taken across all species; the distribution includes values for individual reactions instead of a mean (e.g., a reaction occurring in 7 species has 7 values that are part of the distribution, rather than a single mean as in **Figure 4b**). **d)** Distribution of cluster ratios across all species. As in c, the mean is not taken and individual values are included.

